# CENP-F controls force generation and the dynein-dependent stripping of CENP-E at kinetochores

**DOI:** 10.1101/627380

**Authors:** Philip Auckland, Andrew D. McAinsh

## Abstract

Accurate chromosome segregation demands efficient capture of microtubules by kinetochores and their conversion to stable bi-oriented attachments that can congress and then segregate chromosomes. An early event is the shedding of the outermost fibrous corona layer of the kinetochore following microtubule attachment. Centromere protein F (CENP-F) is part of the corona, contains two microtubule-binding domains and physically associates with dynein motor regulators. Here, we have combined CRISPR gene editing and engineered separation-of-function mutants to define how CENP-F contributes to kinetochore function. We show here that the two microtubule-binding domains make distinct contributions to attachment stability and force generation that are required to minimise errors in anaphase, but are dispensable for congression. We further identify a specialised domain that functions to inhibit the dynein mediated stripping of CENP-E motors. We show how this “dynein-brake” is crucial for ensuring kinetochores contain the right number of CENP-E motors at the right time during mitosis, with loss of brake function delaying congression.

## Introduction

Accurate chromosome segregation during mitosis is dependent upon a centromere associated protein machine called the kinetochore. Kinetochores are built from the hierarchical assembly of two major complexes, the 16-subunit constitutive centromere associated network, which makes multiple contacts with CENP-A nucleosomes and recruits the 10-subunit KMN network (Musacchio and Desai, 2017). Multiple copies of these complexes give rise to the inner and outer kinetochore, and are well established as drivers of microtubule attachment and spindle assembly checkpoint (SAC) signalling (Monda and Cheeseman, 2018; Musacchio, 2015). Metazoan kinetochores also contain a region distal to the outer kinetochore called the ‘fibrous-corona’, named as such due to its appearance on electron micrographs (McEwen, 1993; McEwen et al., 1998). The corona is a highly dynamic structure built from the Rod-Zw10-Zwlich (RZZ) complex, Spindly, Centromere protein (CENP)-F, the molecular motors CENP-E and dynein and the checkpoint protein Mad1 (Maiato et al., 2004). At unattached kinetochores, the corona expands into a crescent-like structure that can even encircle the entire pair of sister chromatids (Hoffman et al., 2001; Magidson et al., 2015; Pereira et al., 2018; Sacristan et al., 2018; Thrower et al., 1996; Wynne and Funabiki, 2015; Wynne and Funabiki, 2016). This expansion is driven by a farnesylation-mediated conformational change in Spindly (Sacristan et al., 2018) and Mps1-dependent phosphorylation of Rod – both of which enable the self-assembly of RZZ-Spindly (RZZS) into a high order meshwork (Pereira et al., 2018; Rodriguez-Rodriguez et al., 2018; Sacristan et al., 2018). This expansion mechanism does not require dynein motor activity, which is instead implicated in corona compaction (Pereira et al., 2018). The expanded corona is thought to provide a large surface area for the initial (lateral) capture of spindle microtubules by CENP-E and dynein motors (Sacristan et al., 2018). As microtubules form end-on kinetochore attachments the corona is disassembled and SAC signalling silenced, both a result, in part, of the dynein-mediated stripping of Mad2 and Rod from kinetochores towards the minus ends of spindle microtubules (Howell et al., 2001; Siller et al., 2005; Wojcik et al., 2001). The contribution of CENP-F to these corona processes is less well understood.

Originally termed mitosin, CENP-F is a large (~360 kDa) coiled-coil protein that dimerises and localises to diverse subcellular locations, including microtubule plus-ends, mitochondria, the nuclear pores and kinetochores (Berto and Doye, 2018; Berto et al., 2018; Kanfer et al., 2015; Rattner et al., 1993). CENP-F contains several non-overlapping functional domains, which include two high affinity microtubule-binding domains (MTBDs), one at either terminus (Feng et al., 2006; Kanfer et al., 2017; Musinipally et al., 2013; Volkov et al., 2015), and binding sites for kinetochore (Bub1; (Berto et al., 2018; Ciossani et al., 2018)), mitochondrial (Miro; (Kanfer et al., 2015; Kanfer et al., 2017; Peterka and Kornmann, 2019)) and nuclear pore (Nup133; (Berto and Doye, 2018; Berto et al., 2018)) adapter proteins. Biochemical analysis of the purified MTBDs revealed that they have distinct microtubule binding characteristics with both able to autonomously track depolymerising microtubule plus-ends in vitro (Volkov et al., 2015). Therefore, the adapter dependent recruitment of CENP-F to subcellular structures allows them to harness microtubule plus-end dynamics to do work. In line with this, a recent work showed that Miro-CENP-F couples mitochondria to dynamic microtubule tips (Kanfer et al., 2015; Kanfer et al., 2017; Peterka and Kornmann, 2019). How these MTBDs, which have a similar microtubule binding affinity to the major attachment factor Ndc80 (Volkov et al., 2015), contribute to kinetochore function in cells is a significant open question.

CENP-F recruitment to kinetochores does not involve the major corona complex RZZS but rather a direct interaction between a defined targeting domain (amino acids 2826-2894) and the kinase domain of Bub1 (Ciossani et al., 2018). Indeed, recruitment of CENP-F to kinetochores is severely compromised in the absence of Bub1 (Ciossani et al., 2018; Currie et al., 2018; Johnson et al., 2004; Liu et al., 2006; Raaijmakers et al., 2018). Furthermore, CENP-F is implicated in the recruitment of the molecular motors dynein (and the Nde1/Ndel1/Lis1 regulators) and CENP-E (Bomont et al., 2005; Faulkner et al., 2000; Simoes et al., 2018; Stehman et al., 2007; Tai et al., 2002; Vergnolle and Taylor, 2007; Yang et al., 2005). Early RNAi studies reported that depletion of CENP-F is associated with severe chromosome congression defects (Bomont et al., 2005; Feng et al., 2006; Holt et al., 2005; Vergnolle and Taylor, 2007; Yang et al., 2005), however, CENP-F knockout mice are viable (Haley et al., 2019) and recent CRISPR knock outs in haploid human cells do not show chromosome alignment defects (Raaijmakers et al., 2018). Defining how CENP-F contributes to chromosome segregation processes thus remains unresolved.

## Results

### CENP-F MTBDs are required for K-K tension and stable microtubule attachment

Due to the pleiotropic phenotypes associated with CENP-F depletion, we reasoned that the two CENP-F microtubule-binding domains could make distinct contributions to kinetochore function. We therefore constructed CENP-F^GFP transgenes (a kind gift from B. Kornmann) that have a GFP inserted at position 1529 between two predicted coiled-coils and either one or both of the MTBDs deleted (see schematic in Figure 1a). First, we confirmed previous work showing that that CENP-E motors were lost (by ~50%) from kinetochores in CENP-F siRNA treated cells (Yang et al., 2005) (Figure 1b,c; n ≥ 2, 150KTs). Expression of all CENP-F mutants restored CENP-E loading (figure 1b,c; n ≥ 2, 150KTs), and both MTBD mutants and WT transgenes bound kinetochores to the same extent (Figure 1c,d n ≥ 3, 515KTs). Next, we quantified the inter-kinetochore distance (K-K) and found that the K-K distance was reduced from 1.49±0.23 *µ*m in control cells to 1.2± 0.22 *µ*m in CENP-F-depleted cells that were transfected with an empty vector (Figure 2e; Supplementary figure 1a; n ≥ 3, 230KTs). Crucially, while transfection with full length CENP-F^GFP rescued this K-K tension to 1.44±0.22*µ*m, cells transfected with CENP-F^GFP∆nMTBD (missing amino terminal MTBD), CENP-F^GFP∆cMTBD (missing the carboxy-terminal MTBD) or CENP-F^GFP∆n+cMTBD (both MTBD deleted) phenocopied the loss of CENP-F with measurements of 1.14±0.23*µ*m, 1.23±0.16*µ*m and 1.23±0.16 *µ*m, respectively (Figure 2e; Supplementary figure 1a; n ≥ 3, 230KTs). Because K-K distance is dependent upon the pulling forces generated by microtubule attachment we then tested the stability of microtubule-kinetochore attachments. To do so, we incubated cells on ice for 10 min prior to fixation and then quantified the kinetochore proximal α-tubulin signal as a readout of microtubule number. In CENP-F siRNA treated cells transfected with an empty vector, the α-tubulin intensities were reduced to 41±28% when compared to control (Figure 1f,g, Supplementary figure 1b n ≥ 3, 270KTs). Transfection with CENPF^GFP or CENP-F^GFP∆cMTBD rescued α-tubulin levels to 88±49% and 85±45%, respectively (Figure 1f,g; Supplementary figure 1b; n ≥ 3, 270KTs). In contrast, cells expressing either CENP-F^GFP∆nMTBD or CENP-F^GFP∆n+cMTBD failed to restore microtubule stability (α-tubulin intensities of 49±26% and 37±18%), demonstrating that the CENP-F N-terminal MTBD is both necessary and sufficient for cold stable microtubule attachment (Figure 1f,g; Supplementary Figure 1b; n ≥ 3, 270KTs). Taken together, these separation-of-function mutants show how microtubule binding by CENP-F is required for stable microtubule-kinetochore attachment and the generation of normal tension across sister kinetochores – but dispensable for the loading of CENP-E motors.

**Figure 1:**
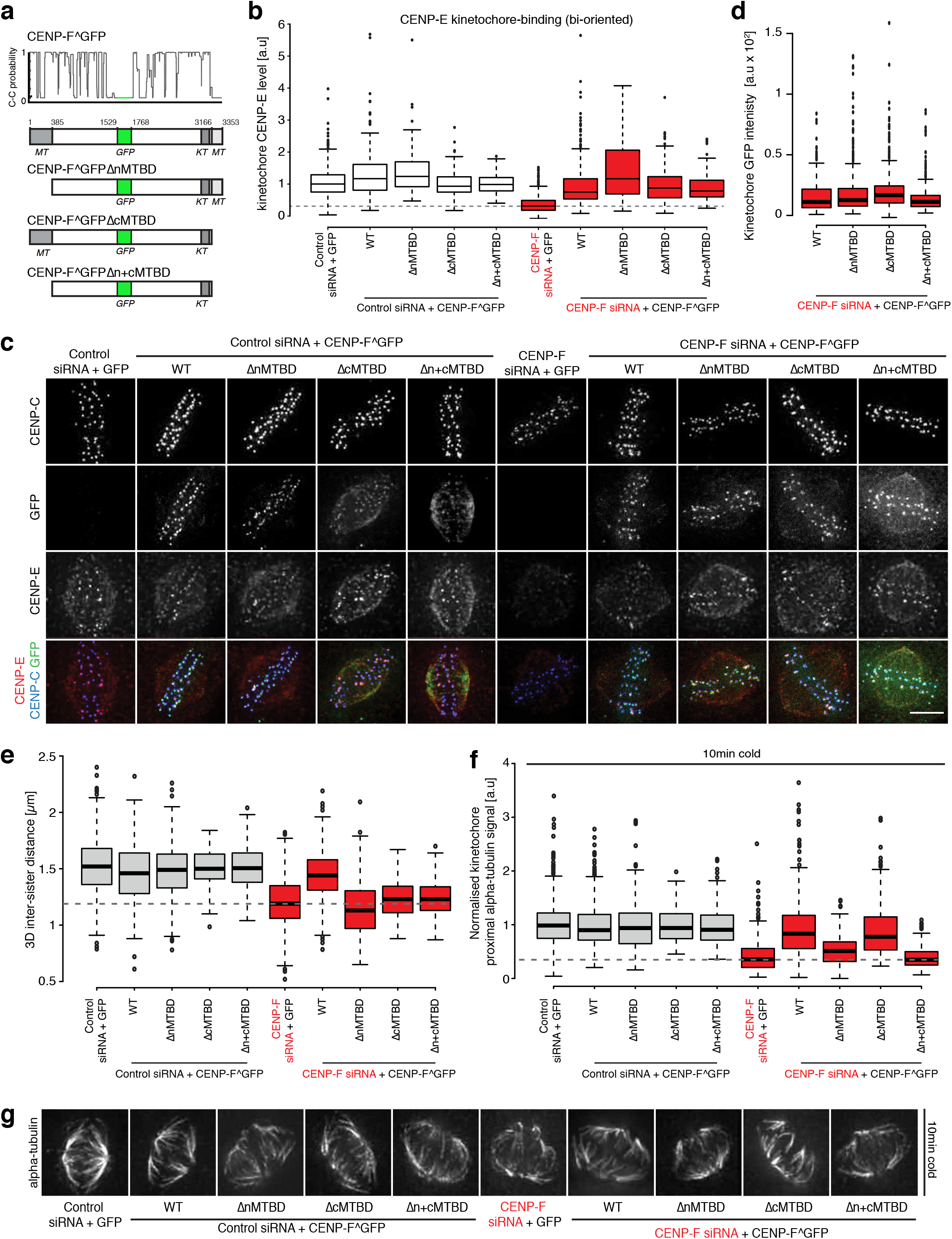
CENP-F MTBDs are required for K-K tension and stable microtubule attachment. **(a)** Cartoon schematic of the CENP-F microtubule-binding mutants and coiled-coil prediction for the CENP-F^GFP construct. **(b)** Quantification of kinetochore CENP-E intensity relative to CENP-C in the rescue experiment depicted in (c). **(c)** Immunofluorescence microscopy images of HeLa-K cells treated with either control or CENP-F siRNA, rescued with wild-type CENP-F or a microtubule binding mutant and stained with DAPI and antibodies against CENP-C and CENP-E. Scale bar 5*µ*m. **(d)** Quantification of kinetochore eGFP intensity in HeLa-K cells transfected with either wild-type CENP-F or the respective microtubule-binding mutants. **(e)** Quantification of the CENP-A based inter-sister distance in the CENP-F rescue experiment depicted in supplementary figure 1a. Dotted grey line indicates the inter-sister distance in cells treated with CENP-F siRNA and rescued with an empty vector. **(f)** Quantification of kinetochore proximal α-tubulin intensity relative to CENP-C in the cold-stable CENP-F rescue experiment depicted in (g) and supplementary figure 1b. Dotted grey line indicates the α-tubulin intensity in cells treated with CENP-F siRNA and rescued with an empty vector. **(g)** Immunofluorescence microscopy images of the cold-stable CENP-F rescue experiment. HeLa-K cells were treated with either control or CENP-F siRNA and rescued with wild-type CENP-F or a microtubule-binding mutant before incubation on ice and fixation. Cells were stained with DAPI and antibodies against CENP-C and α-tubulin. Only the α-tubulin channel is shown. Scale bar 5*µ*m.

**Figure 2:**
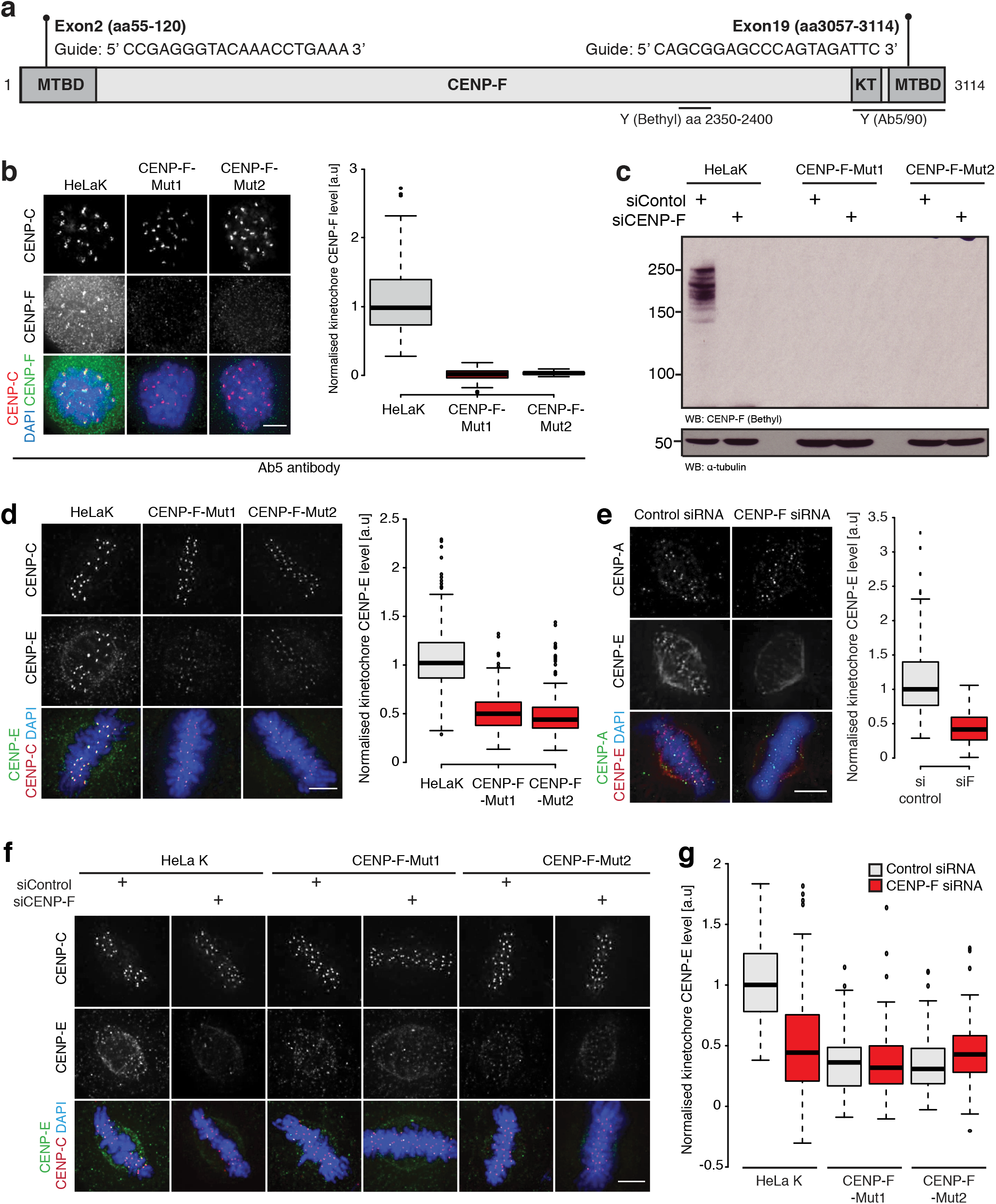
CENP-F CRISPR mutants phenocopy CENP-F RNAi. **(a)** Cartoon schematic of the CRISPR guide targets and antibody epitopes in CENP-F. **(b)** *left* Immunofluorescence microscopy images of HeLa-K, CENP-F-Mut1 and CENP-F-Mut2 cells stained with antibodies against CENP-C and CENP-F (Ab5). Scale bar 5*µ*m. *Right* Quantification of kinetochore CENP-F(Ab5) intensity relative to CENP-C in HeLa-K, CENP-F-Mut1 and CENP-F-Mut2 cells. **(c)** Immunoblot of liquid-N_2_ extracts collected from HeLa-K, CENP-F-Mut1 and CENP-F-Mut2 cells treated with either control or CENP-F siRNA. The membrane was probed with antibodies against CENP-F(Bethyl) and α-tubulin. **(d)** *left* Immunofluorescence microscopy images of HeLa-K, CENP-F-Mut1 and CENP-F-Mut2 cells stained with DAPI and antibodies against CENP-E and CENP-C. Scale bar 5*µ*m. *Right* Quantification of kinetochore CENP-E intensity relative to CENP-C in HeLa-K, CENP-F-Mut1 and CENP-F-Mut2 cells. **(e)** *Left* Immunofluorescence microscopy images of HeLa-K cells treated with either control or CENP-F siRNA and stained with DAPI and antibodies against CENP-E and CENP-A. Scale bar 5*µ*m. *Right* Quantification of kinetochore CENP-E intensity relative to CENP-A in cells treated with either control or CENP-F siRNA. **(f)** Immunofluorescence microscopy images of HeLa-K, CENP-F-Mut1 and CENP-F-Mut2 cells treated with either control or CENP-F siRNA and stained with DAPI and antibodies against CENP-E and CENP-C. Scale bar 5*µ*m. **(g)** Quantification of kinetochore CENP-E level relative to CENP-C in HeLa-K, CENP-F-Mut1 and CENP-F-Mut2 cells treated with control or CENP-F siRNA.

### CENP-F CRISPR mutants phenocopy CENP-F RNAi

Our data shows that CENP-F is required for loading a subset (~50%) of CENP-E to kinetochores (Figure 1b,c). To rule out partial RNAi depletion effects we generated CENP-F mutant cell lines using CRISPR/Cas9 by simultaneously targeting exon 2 and 19 (Figure 2a; (Raaijmakers et al., 2018)) and obtained two independent clonal lines (named CENP-F-Mut1 and CENP-F-Mut2; see Supplementary Figure2d). In both clones we only recovered alleles with premature stop codons and were unable to detect CENP-F protein by quantitative immunofluorescence and western blotting using antibodies against two different C-terminal epitopes (Figure 2b-c, Supplementary Figure 2a,b; n = 1, 100KTs per condition). We note that the Ab5 epitope is partly encoded by exon 19 and could therefore be destroyed by repair of the double strand break. To rule this out, prepared protein extracts from Rpe1 cells stably expressing the CENP-F kinetochore-targeting domain (KTD) fused to Halo-Tag, which is located downstream of the cut site (Figure 2a), and blotted for CENP-F using the Ab5 antibody. As shown in supplementary figure 2c the KTD fragment was clearly detected, confirming that the Ab5 antibody can recognise an epitope downstream of exon19. Finally, we confirmed that long-term passage of these lines did not lead to any (detectable) re-expression of CENP-F, which would be consistent with CENP-F being non-essential in human cells (Supplementary figure 2e-h; n = 1, 100KTs per condition). CENP-E levels on metaphase kinetochores were reduced to 47±18% and 45±19% in CENP-F-Mut1 and CENP-F-Mut2, respectively (Figure 2d; n = 3, 300KTs per condition) – equivalent to that measured in our RNAi experiments (Figure 1b,c, Figure 2e). Moreover, these cells displayed a reduction in K-K distance and had fewer cold-stable microtubules (Supplementary figure 3a-d n = 2, 200KTs per condition). Because trace amounts of functional proteins can remain in CRISPR clones (Meraldi, 2019), we repeated this series of experiments by treating both the CENP-F-Mut1 and CENP-F-Mut2 clones with CENP-F siRNA. In all cases we did not measure any additive phenotypes providing further evidence that the CENP-F Mutants are loss-of-function alleles and that the siRNA is both on-target and efficient at depleting CENP-F (Figure 2f,g; Supplementary figure 3e-h; n = 1, 100KTs per condition).

**Figure 3:**
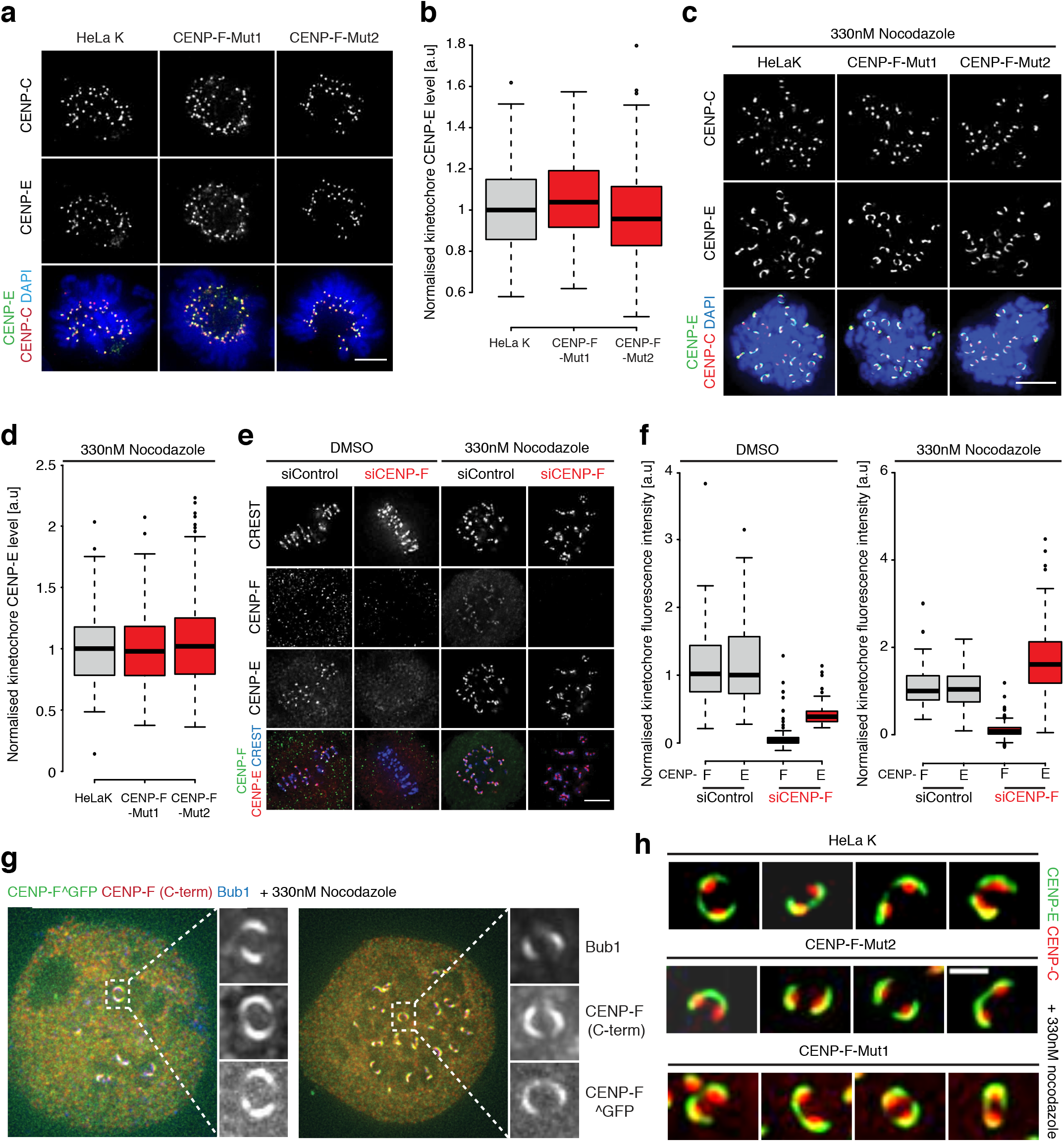
CENP-F controls CENP-E loading in a microtubule-dependent manner. **(a)** Immunofluorescence microscopy images of early prometaphase HeLa-K, CNP-F-Mut1 and CENP-F-Mut2 cells stained with DAPI and antibodies against CENP-C and CENP-E. Scale bar 5*µ*m. **(b)** Quantification of kinetochore CENP-E intensity relative to CENP-C in early prometaphase HeLa-K, CENP-F-Mut1 and CENP-F-Mut2 cells. **(c)** Immunofluorescence microscopy images of HeLa-K, CENP-F-Mut1 and CENP-Mut2 treated with 330nM nocodazole for 16hr and stained with DAPI and antibodies against CENP-C and CENP-E. Scale bar 5*µ*m. **(d)** Quantification of kinetochore CENP-E intensities relative to CENP-C in HeLa-K, CENP-F-Mut1 and CENP-Mut2 treated with 330nM nocodazole for 16hr. **(e)** Immunofluorescence microscopy images of HeLa-K cells treated with control or CENP-F siRNA, incubated with DMSO or 330nM nocodazole for 16hr and stained with DAPI, CREST antisera and antibodies against CENP-E and CENP-F. Scale bar 5*µ*m. The CREST display intensities are not comparable between DMSO and nocodazole conditions. **(f)** Quantification of kinetochore CENP-E and CENP-F levels relative to CREST in HeLa-K cells treated with control or CENP-F siRNA and incubated with DMSO or 330nM nocodazole for 16hr. **(g)** Immunofluorescence microscopy images of HeLa-K cells transfected with CENP-F^GFP, arrested in 330nM nocodazole for 16 hr and stained with antibodies against Bub1 and CENP-Fs C-terminus. Insets show zooms of expanded kinetochores. **(h)** Immunofluorescence microscopy images of CENP-E crescents in HeLa-K, CENP-F-Mut1 and CENP-F-Mut2 cells arrested in 330nM nocodazole for 16hr.

### CENP-F controls CENP-E loading in a microtubule-dependent manner

It is well established that CENP-E motors are displaced from kinetochores as end-on attachments form and bi-orientation is established (Bancroft et al., 2015; Hoffman et al., 2001; Thrower et al., 1996). This raises the possibility that CENP-E kinetochore binding/unbinding maybe sensitive to the presence of attached microtubules. To test this, we quantified the CENP-E intensity at early prometaphase kinetochores in CENP-F-Mut1 and CENP-F-Mut2 cells, a time where sister-pairs lack end-on attachment and are predominantly unattached or laterally attached (Magidson et al., 2011). This revealed that CENP-E was fully loaded to 104±20% and 98±24%, respectively (Figure 3a,b’ n = 2, 200KTs per condition) – even in the absence of CENP-F. To further this observation, we treated parental and CENPF-Mut1 or Mut2 clones with nocodazole and again found that CENP-E motors rebound kinetochores (to 99.7±29.1% and 103.2±35%, respectively; Figure 3c,d; n = 2, 200KTs per condition). We could also confirm this result using CENP-F RNAi treated cells (38±13.8% in DMSO vs. 161.7±100.1% in nocodazole; Figure 3e,f; n = 1, 100KTs per condition). Consistently, both the carboxy-terminus and central regions of CENP-F – like CENP-E -are part of crescent structures at unattached kinetochores (Figure 3g). However, CENP-F is not required for outer kinetochore expansion per se, as CENP-E crescents were clearly visible in both CENP-F-Mut1 and CENP-F-Mut2 clones (Figure 3h). Taken together, these data suggest that CENP-F maintains kinetochore CENP-E levels after end-on microtubule attachment.

### CENP-F functions as a dynein-brake to protect CENP-E from excessive stripping

The reloading of CENP-E to unattached kinetochores suggests that CENP-F controls the motors kinetochore localisation indirectly. One possibility is that CENP-E is stripped by dynein after microtubule attachment and CENP-F regulates this stripping. In line with this, CENP-E is suggested to be a dynein cargo (Howell et al., 2001) and CENP-F recruits the Nde1/Ndel1/Lis1 dynein regulators to kinetochores (Faulkner et al., 2000; Siller et al., 2005; Simoes et al., 2018; Stehman et al., 2007; Tai et al., 2002; Vergnolle and Taylor, 2007). To test this idea, we first sought to directly assay dynein dependent stripping of kinetochore proteins. Previous work used an azide based ATP reduction assay that caused enrichment of cargoes at the spindle poles (Howell et al., 2001), however, this is a non-specific treatment. We reasoned that an alternative approach is the comparison of cells arrested in nocodazole, where cargoes are stably bound, and monastrol, where cargoes are stripped away from the syntelic (end-on) kinetochore-pairs towards the monopole (Figure 4a). This can then be combined with dynein heavy chain (DHC) siRNA to test the motors direct involvement (Figure 4b). To ensure that DHC depletion had no effect on kinetochore-microtubule interaction we stained siControl or siDHC treated cells arrested in monastrol with an antibody against SKAP, a marker for end-on attachment (Schmidt et al., 2010). Quantitation revealed that the SKAP signal was comparable between these conditions, supporting the idea that dynein perturbation does not affect end-on attachment in this setting (Figure 4c,d; n ≥ 2, 190KTs). Next, we aimed to validate the stripping assay and confirm CENP-E as a dynein cargo. To do this, we first compared the kinetochore CENP-E intensities in cells treated with siControl+nocodazole or siControl+monastrol and found that CENP-E levels were reduced to 43.5±35.8% in the siControl+monastrol condition (Figure 4e,f; n ≥ 3, 280KTs). Crucially, DHC depletion in monastrol-arrested cells rescued CENP-E levels to 89±78.3%, confirming that; (1) the nocodazole/monastrol assay can be used to probe dynein-mediated stripping of kinetochore proteins, and (2) CENP-E is a dynein cargo (Figure 4e,f; n ≥ 3, 280KTs). We then tested whether CENP-F regulates dynein stripping by conducting this assay in our CENP-F-Mut1 and CENP-F-Mut2 cell lines. We found that CENP-E levels were further reduced to 9.6±13. 6% and 5.1±7.8% in siControl+monastrol treated CENP-F-Mut1 and CENP-F-Mut2 cells, respectively. In both cases, CENP-E levels were partially rescued by siDHC treatment (Figure 5a-d; n = 2, 200KTs per condition), suggesting that CENP-F negatively regulates dynein activity to prevent excessive CENP-E stripping.

**Figure 4:**
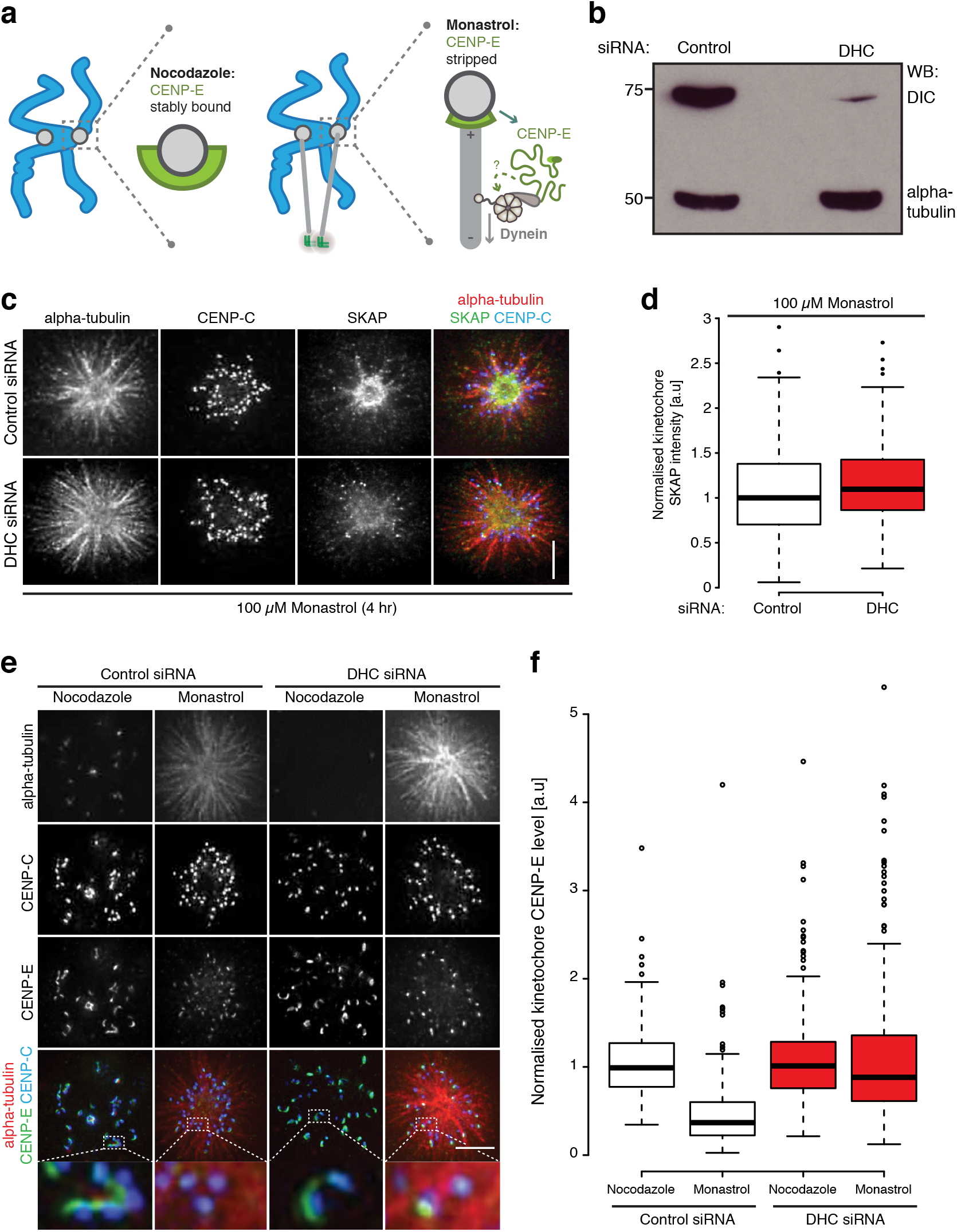
An assay for dynein mediated CENP-E stripping. **(a)** Cartoon schematic of the dynein-stripping assay. Briefly, we compare cells arrested in nocodazole, where dynein cargoes are stably bound, and monastrol, where dynein can transport cargoes away from the syntelic kinetochores towards the monopole. **(b)** Immunoblot of liquid-N_2_ extracts collected from HeLa-K cells treated with control or DHC siRNA. The membrane was probed with antibodies against dynein intermediate chain and α-tubulin. **(c)** Immunofluorescence microscopy images of HeLa-K cells treated with control or DHC siRNA, arrested in 100*µ*M monastrol for 4hr and stained with DAPI and antibodies against SKAP and CENP-C. Scale bar 5*µ*m. **(d)** Quantification of kinetochore SKAP intensities relative to CENP-C in HeLa-K cells treated with control or DHC siRNA and arrested in 100*µ*M monastrol for 4hr. **(e)** Immunofluorescence microscopy images of the dynein stripping assay in HeLa-K cells. Here, cells were treated with control or DHC siRNA, arrested in nocodazole or monastrol and stained with DAPI and antibodies against CENP-E and CENP-C. **(f)** Quantification of the kinetochore CENP-E intensity relative to CENP-C in the dynein-stripping assay depicted in (e).

**Figure 5:**
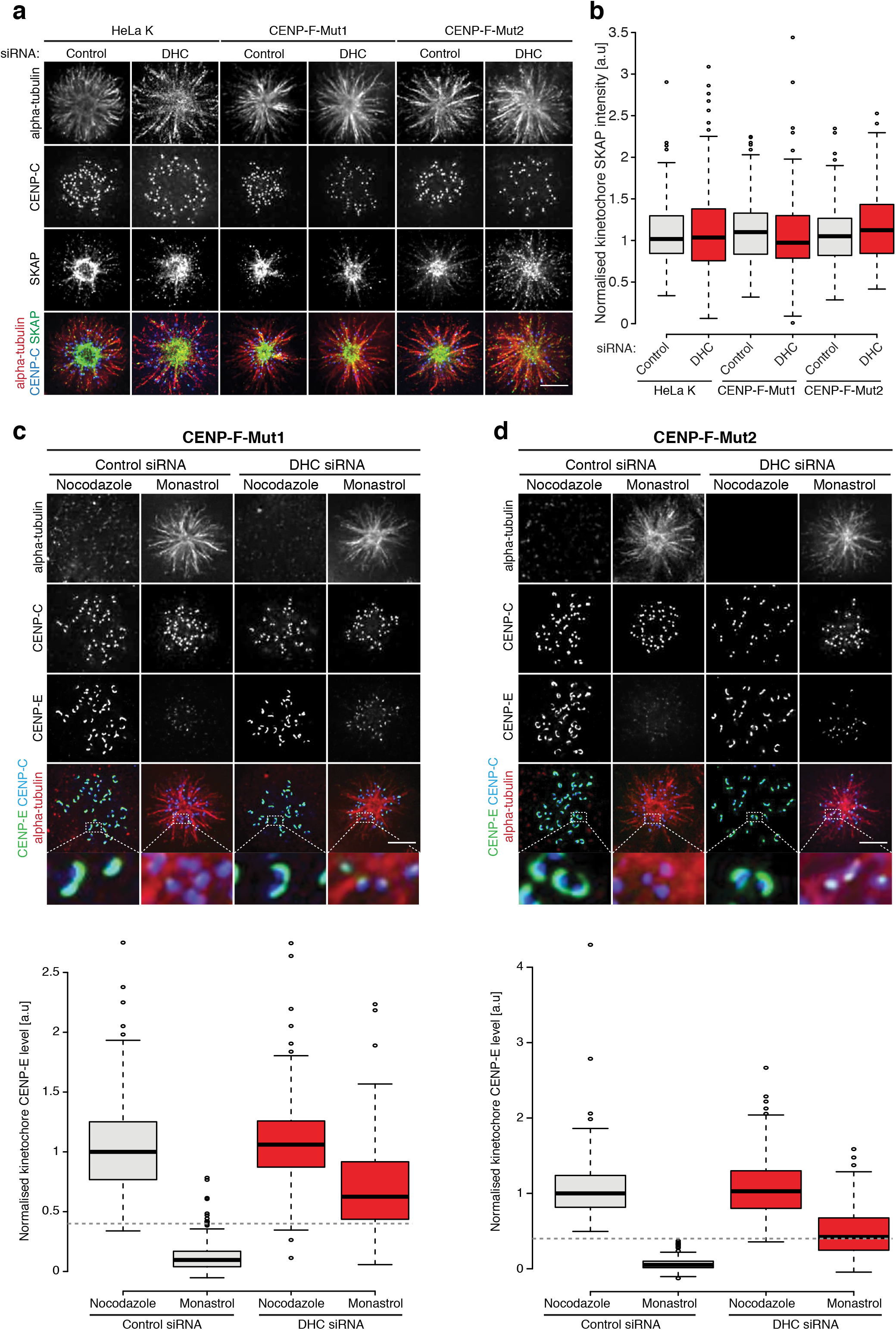
CENP-F functions as a dynein brake that limits CENP-E stripping. **(a)** Immunofluorescence microscopy images of HeLa-K, CENP-F-Mut1 and CENP-F-Mut2 cells treated with control or DHC siRNA, arrested in 100*µ*M monastrol for 4hr and stained with DAPI and antibodies against SKAP and CENP-C. Scale bar 5*µ*m. **(b)** Quantification of kinetochore SKAP intensity relative to CENP-C in HeLa-K, CENP-F-Mut1 and CENP-F-Mut2 cells treated with control or DHC siRNA and arrested in 100*µ*M monastrol for 4hr. **(c)** *Above* Immunofluorescence microscopy images of the dynein stripping assay in CENP-F-Mut1 cells. Here, cells were treated with control or DHC siRNA, arrested in nocodazole or monastrol and stained with DAPI and antibodies against CENP-E and CENP-C. Scale bar 5*µ*m. *Below* Quantification of the kinetochore CENP-E intensity relative to CENP-C in the dynein-stripping assay depicted above. Dotted grey line indicates the level of CENP-E stripping in HeLa-K cells treated with control siRNA and arrested in monastrol, see Figure 6f. Note that CENP-E levels are further reduced, indicating an upregulation of dynein mediated stripping. **(d)** *Above* Immunofluorescence microscopy images of the dynein stripping assay in CENP-F-Mut2 cells. Here, cells were treated with control or DHC siRNA, arrested in nocodazole or monastrol and stained with DAPI and antibodies against CENP-E and CENP-C. Scale bar 5*µ*m. *Below* Quantification of the kinetochore CENP-E intensity relative to CENP-C in the dynein-stripping assay depicted above. Dotted grey line indicates the level of CENP-E stripping in HeLa-K cells treated with control siRNA and arrested in monastrol, see Figure 6f. Note that CENP-E levels are further reduced, indicating an upregulation of dynein mediated stripping.

To gain mechanistic insight into how CENP-F regulates kinetochore-associated dynein, we created two eGFP tagged truncations of CENP-F (Figure 6a). The first contains the binding site for Nde1/Ndel1, which recruits the dynein regulator Lis1 to kinetochores, an adjacent coiled-coil region and the kinetochore-targeting domain (KTD) (eGFP-CENP-F(2021-2901)) (Figure 6a) (Stehman et al., 2007; Vergnolle and Taylor, 2007; Zhu et al., 1995). The second lacks the Nde1/Ndel1 binding site and contains only the coiled-coil region and KTD (eGFP-CENP-F(2351-2901)) (Figure 6a). Both transgenes were equally loaded to kinetochores, ruling out the expression level as a phenotypic driver (Figure 6b,c; n ≥ 4, 681KTs). We then tested if transient expression of these fragments in CENP-F-Mut1 and CENP-F-Mut2 cells altered CENP-E levels at metaphase kinetochores. This revealed that eGFP-CENP-F(2021-2901) could rescue CENP-E levels from ~30% to 71.4±36.6% and 67.3±39% in CENP-F-Mut1 and CENP-F-Mut2 cells, respectively (Figure 6b,d; n ≥ 3, 280KTs). Crucially, expression of eGFP-CENP-F(2351-2901) phenocopied transfection with an empty vector (Figure 6b,d; n ≥ 3, 280KTs). Taken together, these data show that CENP-F regulates the dynein mediated stripping of CENP-E, and this is likely via direct interaction with Nde1/Ndel1.

**Figure 6:**
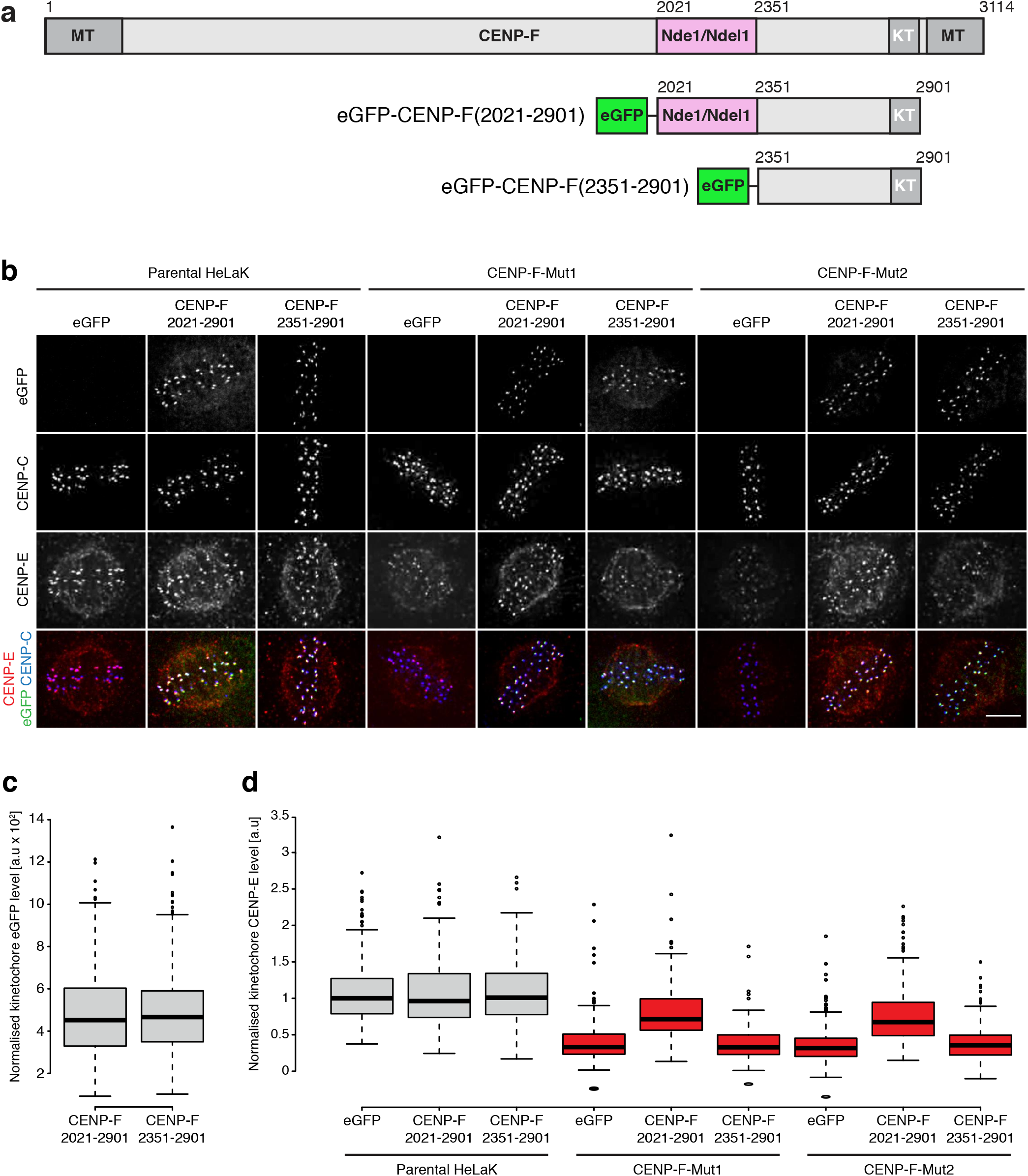
The dynein brake requires the interaction of CENP-F with Nde1/Ndel1. **(a)** Cartoon schematic of the CENP-F Nde1/Ndel1 binding mutants. **(b)** Immunofluorescence microscopy images of HeLa-K, CENP-F-Mut1 and CENP-F-Mut2 cells transfected with either an empty vector, eGFP-CENP-F(2021-2901) or eGFP-CENP-F(2351-2901) and stained with DAPI and antibodies against CENP-E and CENP-C. Scale bar 5*µ*m. **(c)** Quantification of kinetochore eGFP intensities minus background in cells expressing eGFP-CENP-F(2021-2901) or eGFP-CENP-F(2351-2901). **(d)** Quantification of kinetochore CENP-E intensities relative to CENP-C in HeLa-K, CENP-F-Mut1 and CENP-F-Mut2 cells transfected with either an empty vector, eGFP-CENP-F(2021-2901) or eGFP-CENP-F(2351-2901).

### Distinct functional contributions of CENP-F dynein-brake and MT binding activities

Having established cell lines with undetectable CENP-F we sought to understand the functional importance of CENP-F in human cells. To do so, HeLa-K, CENP-F-Mut1 and CENP-F-Mut2 cells were labelled with SiR-DNA and imaged every 3 min for 12 hr and the timings of chromosome congression and anaphase onset determined. Both CENP-F-Mut1 and CENP-Mut2 cells progressed though mitosis and initiated anaphase, albeit with a mild ~6 min delay in chromosome congression (10% unaligned at T=24 min in CENP-F-Mut1 and CENP-F-Mut2 cells compared to 1% in control cells; Figure 7a,b, Supplementary figure 4; n ≥ 3, 300 cells). We note that inter-kinetochore tension is reduced in these cell lines thus showing how normal tension is not necessary for checkpoint satisfaction. Previous work showed how tensionless kinetochores progress normally into anaphase but have an increased error rate that is associated with slower anaphase chromosome movements (Dudka et al., 2018). By filming anaphase HeLa cells expressing eGFP-CENP-A and treated with control or CENP-F siRNA we found that early anaphase kinetochore velocity was reduced from 3.14±1.2 *µ*m min^−1^ in controls cells to 2.34±0.8 *µ*m min^−1^ in CENP-F-depleted cells (Figure 7c,e; n ≥ 100KTs per condition). Moreover, these cells displayed an increased rate of anaphase errors (Figure 7d,f; n ≥ 3, 300 cells).). Taken together, these data show that CENP-F is important for proper anaphase kinetochore dynamics and error-free chromosome segregation, consistent with a role in kinetochore-based force generation.

**Figure 7:**
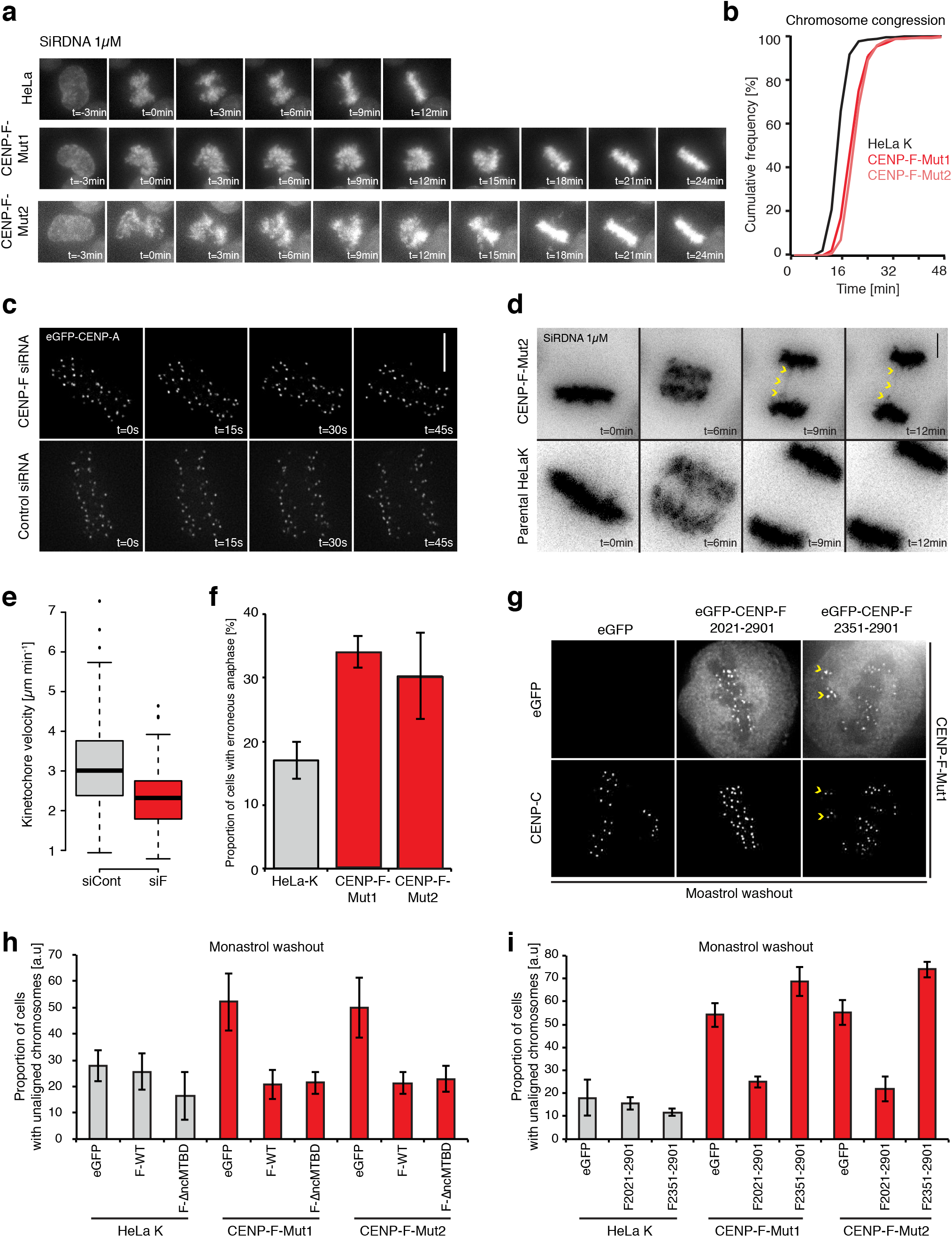
Distinct functional contributions of CENP-F Dynein-brake and MT binding activities. **(a)** Movie stills of HeLa-K, CENP-F-Mut1 and CENP-F-Mut2 cells labelled with 1*µ*M SirDNA progressing from nuclear envelope breakdown to metaphase. **(b)** Cumulative frequency plot of congression timings for HeLa-K, CENP-F-Mut1 and CENP-F-Mut2 cells. **(c)** Stills of early anaphase HeLa cells expressing eGFP-CENP-A and treated with control or CENP-F siRNA. Scale bar 5*µ*m. **(d)** Stills of live HeLa-K and CENP-F-Mut2 cells labelled with 1*µ*M SirDNA progressing through anaphase. Yellow arrows depict a lagging chromosome. Scale bar 5*µ*m. **(e)** Quantification of early anaphase kinetochore velocities in live HeLa cells expressing eGFP-CENP-A and treated with control or CENP-F siRNA. Velocity measurements were taken from tracks of persistent movement that lasted at least 3 time frames. **(f)** Quantification of anaphase error rates in live HeLa-K, CENP-F-Mut1 and CENP-F-Mut2 cells labelled with 1*µ*M SirDNA. Error bars ±SD. **(g)** Immunofluorescence microscopy images of CENP-F-Mut1 cells transfected with eGFP, eGFP-CENP-F(2021-2901) or eGFP-CENP-F(2351-2901) 1 hr post monastrol washout and stained with an antibody against CENP-C. **(h)** The proportion of HeLa-K, CENP-F-Mut1 and CENP-F-Mut2 cells transfected with eGFP, CENP-F^GFP or CENP-F^GFP∆n+cMTBD with unaligned chromosomes 1 hr after monastrol release. **(i)** The proportion of HeLa-K, CENP-F-Mut1 and CENP-F-Mut2 cells transfected with eGFP, eGFP-CENP-F(2021-2901) or eGFP-CENP-F(2351-2901) with unaligned chromosomes 1 hr after monastrol release.

Our data show that CENP-F has two functional modules, one that controls microtubule binding and one that regulates dynein activity. A key question pertains to how these activities are integrated to ensure accurate mitotic progression. To test this, we needed to use to a fixed cell assay in which we could phenotype cells that had WT or mutant CENP-F-GFP-localised to kinetochores. To do this, we arrested and released cells from a monastrol block – this traps kinetochores in a syntelic state around a monopole and requires successful error correction and then chromosome congression to the spindle equator. The fraction of cells with unaligned kinetochores (failed error correction and/or congression) was elevated in both CENP-F-Mut1 and CENP-Mut2 cells (with ~60% of CENP-F-Mut1 and CENP-F-Mut2 cells with unaligned chromosomes *vs.* ~20% in control cells after 1 hr release from monastrol block, Figure 7g-i; Supplementary figure 5a,b; n ≥ 3, 100 cells per condition). This phenotype was clearly more penetrant than that measured in unperturbed cells by live imaging (Figure 7a,b; see discussion). The alignment defect could be rescued by transfection of the WT CENP-F transgene, and surprisingly, with transfection of a CENP-F construct that lacks both MT binding domains (Figure 7h; Supplementary figure 5a; n ≥ 3, 100 cells per condition). This result suggests that the loss of K-K tension and attachment stability following loss of the microtubule binding domains does not impact error correction and congression. Next, we tested whether the dynein-brake alone is sufficient to rescue the monastrol-release phenotype. To do this we transfected HeLa-K, CENP-F-Mut1 and CENP-F-Mut2 cells with eGFP-CENP-F(2021-2901), which contains only the kinetochore targeting and dynein-braking domains. This CENP-F truncation was able to rescue the chromosome congression phenotype (~25% of CENP-F-Mut1 and CENP-F-Mut2 cells with unaligned chromosomes compared to ~19% in control cells after 1hr release from monastrol block), while a control construct that lacks the dynein brake (eGFP-CENP-F(2351-2901) was not able to rescue (Figure 7h,i; Supplementary figure 5b; n ≥ 3, 100 cells per condition). These data show how CENP-F modulation of dynein cargos (including CENP-E) are crucial for efficient conversion of syntelic attachments into aligned and bi-orientated sister kinetochores.

## Discussion

In this work we have defined for the first time how the CENP-F microtubule binding domains contribute to kinetochore activity and identified a novel functional domain that modulates dynein motor activity at kinetochores. We have named this the dynein-brake.

Our data shows how both MTBDs are required for the generation of centromeric tension, while the amino-terminal MTBD also contributes to the stability of kinetochore-microtubule attachments (Figure 8a). This may reflect subtle differences in their biochemical properties, as the amino-terminal MTBD preferentially binds curved microtubule structures while the carboxy-terminal MTBD has a higher affinity for straight lattice configurations (Volkov et al., 2015). We propose the former stabilises depolymerising microtubule plus-ends and thus protects K-fibres from cold induced disassembly. In contrast, both MTBD-microtubule interactions support force generation, which suggests that kinetochores utilise diffusion-based and curved-protofilament coupled mechanisms to produce tension. It is well established that the latter is mediated by the Ska complex (Schmidt et al., 2012), which is crucial for providing load-bearing capability to kinetochores *in vivo* (Auckland et al., 2017) and *in vitro* (Helgeson et al., 2018). However, the CENP-F MTBDs do not appear to contribute as they are dispensable for chromosome congression. Crucially, we found no additive defects when the Ska complex is depleted in CENP-F mutant cells (data not shown) suggesting that this is not due to redundancy. Instead, our evidence supports the idea that CENP-F force generation is important for normal anaphase chromosome speeds and the resolution of merotelic attachments. This is consistent with recent work also showing how low centromere tension does not impact pre-anaphase events but leads to elevated frequency of lagging chromosomes (Dudka et al., 2018).

**Figure 8:**
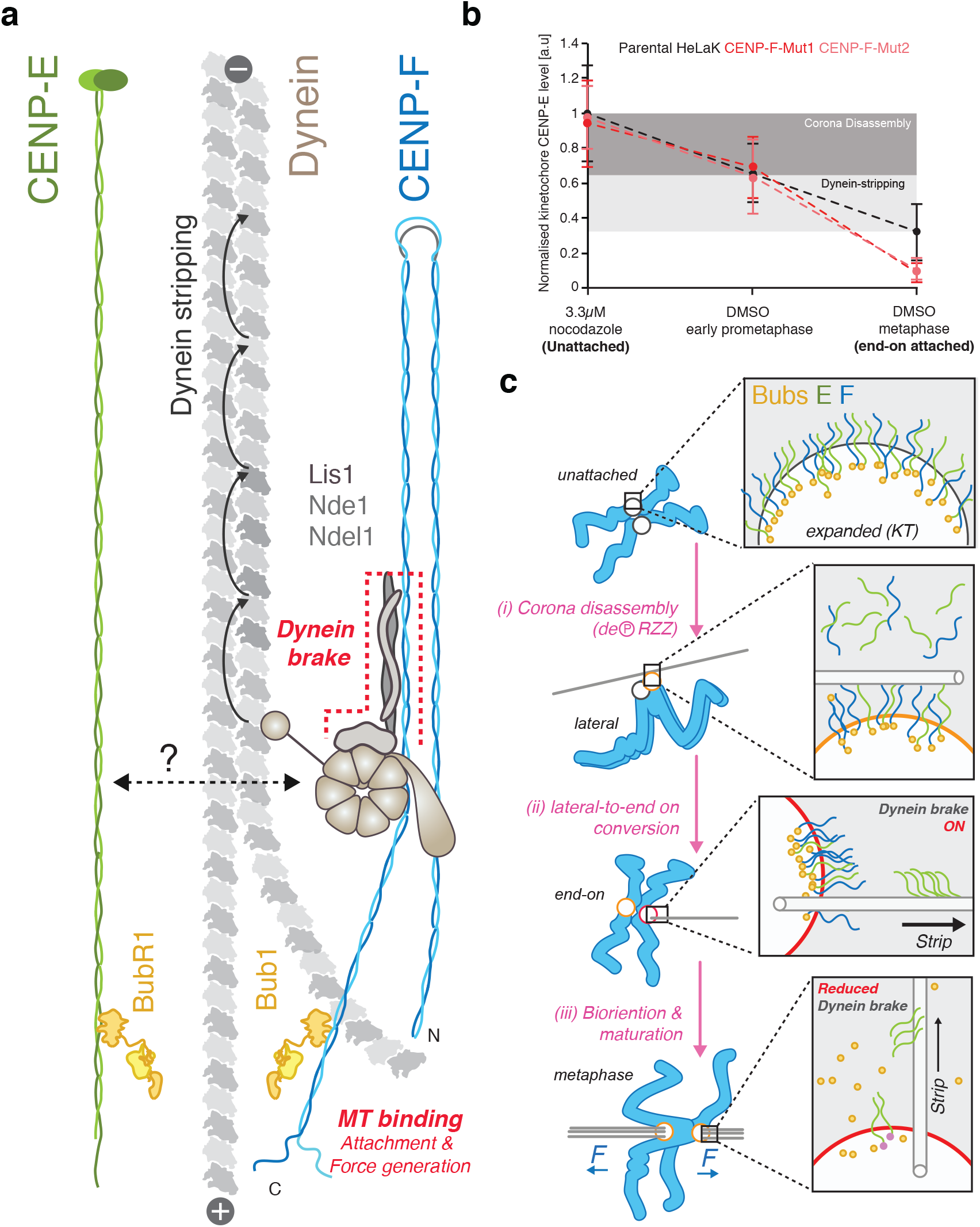
The major functions of CENP-F at kinetochores. **(a)** Schematic (to scale) illustrating major CENP-F functions and molecular interactions at the kinetochore. The two-microtubule binding domains, located at either terminus, bind distinct tubulin sites to generate K-K tension and stabilise end-on microtubule attachments. The C-terminal ~300 amino acid domain acts as a dynein-brake, which limits the rate of CENP-E stripping through physical interaction with known dynein regulators Nde1/Ndel1/Lis1 (Stehman et al., 2007; Vergnolle and Taylor, 2007). **(b)** Summary of CENP-E kinetochore loading in nocodazole, early prometaphase and metaphase in parental, CENP-F-Mut1 and CENP-F-Mut2 HeLa cells. We suggest that the unloading of CENP-E is biphasic: As kinetochores transition from an unattached to laterally attached state the corona is contracted, this unloads ~35% of the motor pool and is unaffected by loss of CENP-F (dark grey region, black and red dotted lines). As these lateral attachments are converted to end-on and subsequently bi-oriented attachments, dynein strips away a further ~50% of the remaining CENP-E motors (light grey region, black dotted line). The loss of the CENP-F dynein-brake exacerbates this stripping, as CENP-F-Mut1 and CENP-F-Mut2 cells display a ~85% reduction in kinetochore CENP-E between prometaphase and metaphase (light grey region, red dotted lines). **(c)** Model of CENP-E dynamics as a function of attachment state during a normal mitosis. This highlights the biphasic nature of CENP-E removal, first by corona compaction and then by dynein mediated stripping as end-on attachments form and mature. The system eventually equilibrates due to the action of the CENP-F dynein-brake, which ensures sufficient motors remain associated with kinetochores to facilitate timely chromosome congression (see discussion for details).

Both the carboxy-terminus and central region of CENP-F are located in the corona of unattached kinetochores along with the targeting subunit Bub1 (Figure 8a; Figure 3g). We find that CENP-F does not support corona formation nor does it recruit CENP-E to kinetochores through a direct physical interaction. Instead, our data indicate that the dynein-brake regulates the rate of corona contraction by inhibiting the activity of dynein motors – which would strip cargos (including CENP-E) from kinetochores. Figure 8b and supplementary figure 5c synthesises our measurements of CENP-E kinetochore binding: the motor is displaced from kinetochores as they progress from an unattached state to prometaphase and then to metaphase. Initial loss of CENP-E (corona compaction; dark grey zone) is independent of CENP-F activity, while the second phase of removal is dynein-dependent and under the control of CENP-F (light grey zone). Our working model (Figure 8c) is that as kinetochores form lateral attachments the corona is compacted and this is triggered by dephosphorylation of the RZZ complex (Step I; (Pereira et al., 2018; Sacristan et al., 2018). As end-on attachments form (Step ii) dynein motor complexes strip cargos (including CENP-E) from the kinetochore. The CENP-F brake puts a limit on this process to ensure a pool of CENP-E motors (and potentially other cargos) remain on kinetochores. As sister kinetochores bi-orientate and align (Step iii) Bub1 is lost from kinetochores, leading to a reduction in CENP-F and the associated brake activity.

We propose that this brake function is mediated through the reported physical interaction with Nde1/Ndel1 (Figure 8c; (Vergnolle and Taylor, 2007), which in turn, recruits the dynein regulator Lis1 to kinetochores (Stehman et al., 2007). Lis1 is a highly conserved and well-established dynein regulator that is mutated in the neurodevelopmental disease Type-1 lissencephaly (Reiner and Coquelle, 2005). Lis1 binds directly to the dynein motor domain and differentially regulates its transport in response to AAA3 nucleotide state (DeSantis et al., 2017; Huang et al., 2012; Toropova et al., 2014). This context-dependent regulation explains how perturbation of Lis1 in cells can both augment and reduce cargo transport at distinct subcellular locations (Dix et al., 2013; Klinman and Holzbaur, 2015; Moughamian et al., 2013; Pandey and Smith, 2011; Shao et al., 2013; Vagnoni et al., 2016; Yi et al., 2011). Our data supports the idea that CENP-F/Ndel1/Nde1/Lis1 at human kinetochores favours a high-binding low-velocity dynein motor state that attenuates CENP-E stripping after end-on attachment (DeSantis et al., 2017). This idea will need testing with in vitro reconstitution experiments. How CENP-E physically associates with Dynein complexes is a cargo is also unknown.

Our experiments are beginning to reveal how the Bub1-CENP-F/Ndel1/Nde1/Lis1 axis contributes to early mitosis in humans (independently of microtubule binding activities). CENP-F-Mut1 and CENP-F-Mut2 cells both have a mild congression delays that can be exacerbated by arrest and release from monastrol. This assay demands that syntelic kinetochores about a monopole are detached from microtubules (through error correction), form lateral attachments that are converted to end-on and ultimately bi-oriented attachments. Because the brake domain can rescue these congression defects we propose that Bub1-CENP-F plays a crucial role in ensuring kinetochores retain the optimal number of CENP-E motors to transition between these key attachment states. The next challenges will be to understand (i) how the CENP-F dynein-brake activity is coordinated with RZZ-Spindly, which brings active dynein to the corona, and (ii) how CENP-F molecules can expand within the self-assembling RZZS corona meshwork.

## Material and methods

### Cell culture, siRNA transfection and drug treatments

HeLa-Kyoto (K), CENP-F-Mut1 and CENP-F-Mut2 cells were grown in a humidified incubator at 37°C and 5% CO_2_ in DMEM (Gibco) containing 10% FCS (Sigma), 100U/ml penicillin and 100*µ*g/ml streptomycin (Gibco). This was supplemented with 0.1*µ*g/ml puromycin (Invitrogen) for the maintenance of the eGFP-CENP-A cell line. The hTERT-RPE1-CENP-F(KTD) cell line was maintained in DMEM/F-12 medium containing 10% FCS, 2.3 g/l sodium bicarbonate, 100 U/ml penicillin, and 100 *µ*g/ml streptomycin. siRNA oligonucleotides (53 nM) were transfected using oligofectamine (Invitrogen) according to the manufacturers guidelines and analysed at 48hr. The following sequences were used: control, 5’-GGACCUGGAGGUCUGCUGU-3’, CENP-F, 5’-AAGAGAAGACCCCAAGUCAUC-3’ (Holt et al., 2005), CENP-E 5’-ACUCUUACUGCUCUCCAGU-3’ (Bancroft et al., 2015) and DHC 5’-GGAUCAAACAUGACGGAAU-3’. For drug treatments, cells were treated with 330nM nocodazole for 16hr or 100 *µ*M monastrol for 4hr before fixation. For monastrol release, cells were arrested for 4hr, washed three times in warm DMEM and incubated for 1hr before fixation. For overnight imaging, cells were incubated with 1 *µ*M SiRDNA (Spirochrome) for 30 min prior to imaging.

### Plasmid construction and siRNA rescue experiments

We note that Uniprot and Ensembl previously reported conflicting CENP-F lengths of 3200 and 3114 amino acids, respectively (this has now been corrected to 3114). This resulted from a 96 amino acid insertion at position 1515 between two major coiled-coil regions. In the present study, all C-terminal domain positions have been adjusted for this insertion. To generate an siRNA protected version of CENP-F^GFP for transient expression in cells, full length CENP-F^GFP was excised from pcDNA5/frt/to using BamHI and NotI sites and cloned into pcDNA3.1+ creating pcDNA3.1+CENP-F^GFP (pMC619). A short DNA fragment containing the siRNA protected sequence (5’GAAAAAACGCCTAGCCACC3’) was synthesised with flanking BamHI and AgeI sites (pMC618, GeneArt) and cloned into pcDNA3.1+CENP-F^GFP to create pcDNA3.1+CENP-F^GFP-RIP (pMC621). The pcDNA3.1+CENP-F^GFP-RIP plasmid was then used to create all microtubule binding mutants. For deletion of the N-terminal MTBD, a fragment of CENP-F from the terminus of the MTBD to an internal Age1 site was amplified by PCR with a flanking 5’ BamHI site. This was cloned into pcDNA3.1+CENP-F^GFP-RIP using BamHI and AgeI sites, creating pcDNA3.1+CENP-F^GFP∆nMTBD (pMC620). For deletion of the C-terminal MTBD, a fragment of CENP-F from an internal SfiI site to the start of the MTBD was amplified by PCR with a flanking 3’ NotI site. This was cloned into pcDNA3.1+CENP-F^GFP-RIP using SfiI and NotI sites to create pcDNA3.1+CENP-F^GFP-RIP∆cMTBD (pMC626). To generate pcDNA3.1+CENP-F^GFP∆n+cMTBD (pMC627) the above cloning was performed sequentially. For siRNA rescue experiments HeLa K cells were grown on 22 mm 1.5 coverglass for 24 hr before treatment with either control or CENP-F siRNA for 24 hr in 1.5 ml MEM. Cells were then transfected with 2 *µ*g of maxi-prep plasmid DNA using FugeneHD at 1:4 according to the manufacturers guidelines and incubated for a further 48 hr in 2 ml DMEM. To generate the Nde1/Ndel1 binding mutants, either CENP-F(2021-2901) or CENP-F(2351-2901) was amplified by PCR with flanking Kpn1 and BamHI sites and cloned into peGFP-C1, creating eGFP-CENP-F(2021-2901) (pMC651) and eGFP-CENP-F(2351-2901) (pMC652), respectively. For expression in cells, 2 **µ**g of plasmid DNA was transfected using FugeneHD at 1:4 and incubated for 48 hr in 2 ml DMEM.

### CRISPR-Cas9

To target CENP-F exons 2 and 19, the guides 5’-CCGAGGGTACAAACCTGAAA-3’ (exon2) and 5’-CAGCGGAGCCCAGTAGATTC-3’ (exon19) (Raaijmakers et al., 2018) were cloned into the human codon optimised SpCas9 and chimeric guide expression plasmid (pX330, Addgene) using BbsI as previously described (Ran et al., 2013). To generate CENP-F-Mut1 and CENP-F-Mut2 cell lines, HeLa-K cells grown in a 33 mm dish were simultaneously transfected with 1 *µ*g of each Cas9-guide plasmid using FugeneHD at 1:4 according to the manufacturers guidelines and incubated for 48 hr. Cells were diluted 10-fold and plated on 15 cm dishes in DMEM without selection and grown for ~2 weeks. Single colonies were picked using typsin soaked cloning disks, amplified and screened by immunofluorescence. The exon2 and exon19 alleles in CENP-F mutant clones were amplified by PCR with flanking Kpn1 and AgeI sites and cloned into peGFP-N1 for sequencing. A summary of the sequencing data can be found in Supplementary figure 2d. At least 10 alleles were sequenced per cut site.

### Immunofluorescence microscopy

Cells were fixed in 20 mM Pipes pH 6.8, 10 mM EGTA, 1 mM MgCl2, 0.2% Triton X-100, and 4% formaldehyde (PTEMF). For the cold stable assay, cells were incubated in ice cold DMEM for 10 min prior to fixation. Cells were washed three times in PBS before blocking in PBS+3% BSA for 30 min and incubation with primary antibodies for 1hr at room temperature: anti-CENP-C (1/2000, Guinea Pig, MBL), anti-CENP-E (1/1000, Rabbit, Meraldi lab), anti-CENP-F(Ab5) (1/400, Rabbit, Abcam), anti-CENP-F(Ab90) (1/400, Mouse, Abcam), anti-CENP-A(1/500, Mouse, Abcam), anti-α-tubulin (1/1000, Mouse, Sigma), anti-SKAP (1/400, Rabbit, Atlas Antibodies), anti-Bub1(Ab54893) (1/200, Mouse, Abcam) and CREST antisera (1/250, human, Antibodies Inc). Cells were washed three times with PBS and incubated with alexa-fluour conjugated secondary antibodies at 1/500 for 1 hr at room temperature. Cells were mounted and imaged in Vectashield (Vector Laboratories). 3D image stacks were acquired using a 100× oil NA 1.4 objective on an Olympus Deltavision Elite microscope (Applied Precision Ltd.) equipped with a DAPI, FITC, Rhodamine, or Texas Red and Cy5 filter set, solid-state light source, and a CoolSNAP HQ2 camera (Roper Technologies) at 37°C. Stacks were deconvolved using SoftWorx and fluorescence intensity measurements were made manually after background subtraction and normalization.

### Live-cell imaging

To film kinetochore fates, HeLa-K eGFP-CENP-A–expressing cells were seeded in FluoroDishes (World Precision) and imaged in DMEM supplemented with 10% FCS, 100 U/ml penicillin, 100 *µ*g/ml streptomycin, and 0.1 *µ*g/ml puromycin. 3D image stacks (25 × 0.5-*µ*m z-sections) were acquired every 7.5 s using a 100× oil NA 1.4 objective on an Olympus Deltavision Elite microscope equipped with a eGFP and mCherry filter set, Quad-mCherry dichroic mirror, solid-state light source, and CoolSNAP HQ2 camera. Environment was tightly controlled at 37°C and 5% CO_2_ using a stage-top incubator (Tokai Hit) and a weather station (Precision Control). Image stacks were deconvolved using SoftWorx (Applied Precision Ltd.), and kinetochore fates were determined manually. Measurements of kinetochore velocity were taken manually from tracks of persistent movement that lasted at least three time frames. For overnight imaging, cells were seeded in FluroDishes (World Precision) and imaged in DMEM supplemented with 10% FCS, 100 U/ml penicillin and 100 *µ*g/ml streptomycin. 3D images stacks (7 × 2 *µ*m z-sections) were acquired every 3 min for 12 hr on the Deltavision system described above.

### Immunoblotting

Protein extracts were prepared by liquid nitrogen grinding. Briefly, cells arrested in 330nM nocodazole for 16hr were harvested from a 15cm dish and resuspended in 1.5× pellet volumes of H-100 buffer (containing 50mM HEPES pH7.9, 1mM EDTA, 100mM KCl, 10% glycerol, 1mM MgCl_2_ and a complete protease inhibitor tablet (Roche)). Cells were ground in liquid N_2_ with a precooled mortar and pestle. The ground cell extract was collected and spun at 14,000rpm for 30min at 4 degrees and the soluble fraction collected. Protein concentration was determined by Bradford. 30*µ*g of extract was boiled in LDS sample buffer + reducing agent (NuPage) for 10min and separated on a 4-12% Bis-Tris gel (NuPage) in MOPS (NuPage). Proteins were wet transferred to a nitrocellulose membrane before blocking in 5% milk TBST for 1hr at room temperature. Primary antibodies were incubated overnight at 4°C in 2% milk TBST: anti-CENP-F(Ab5) (1/1500, Rabbit, Abcam), anti-CENP-F(A301-611A) (1/5000, Rabbit, Bethyl), anti-DIC (1/1000, Mouse, EMD Millipore) and anti-α-tubulin (1/10,000, Mouse, Sigma). Membranes were washed three times in TBST before incubation with HRP-conjugated secondary antibodies for 1hr at room temperature in 2% milk TBST (1/10,000, GE Healthcare).

## Acknowledgements

We thank B. Kornmann (Oxford) for the CENP-F^GFP plasmid and E. Roscioli (Warwick) for generation of the hTERT-RPE1-CENP-F(KTD) cell line. We thank J. A. Millar and E. Roscioli for critical reading of the manuscript. A.D. McAinsh is supported by a Wellcome Trust Senior Investigator Award (grant 106151/Z/14/Z) and a Royal Society Wolfson Research Merit Award (grant WM150020).

## Author contributions

Project conception, planning, data interpretation and manuscript preparation was carried out by A.D.M and P.A. All experiments were performed and analysed by P.A.

---

**Supplementary figure 1.**
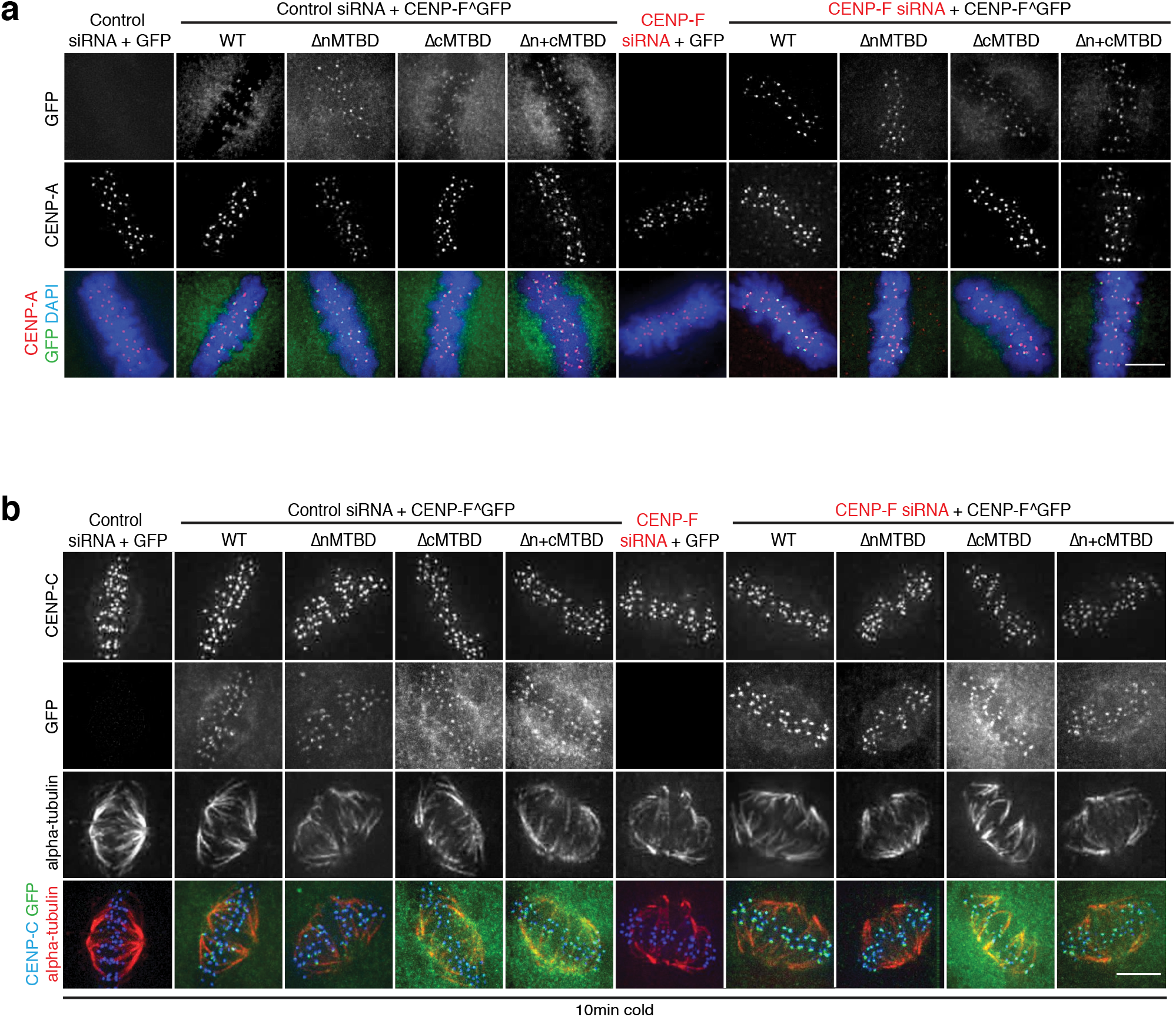
**(a)** Immunofluorescence microscopy images of the CENP-F MTBD K-K distance rescue experiment. HeLa-K cells were treated with control or CENP-F siRNA and rescued with an empty vector, CENP-F^GFPΔnMTBD, CENP-F^GFPΔcMTBD or CENP-F^GFPΔn+cMTBD before being stained with DAPI and an antibody against CENP-A. Scale bar 5μm. **(b)** Immunofluorescence microscopy images of the CENP-F MTBD cold-stable rescue experiment. HeLa-K cells were treated with control or CENP-F siRNA and rescued with an empty vector, CENP-F^GFPΔnMTBD, CENP-F^GFPΔcMTBD or CENP-F^GFPΔn+cMTBD before being incubated on ice for 10min and stained with DAPI and antibodies against CENPC and α-tubulin. Scale bar 5μm.

**Supplementary figure 2.**
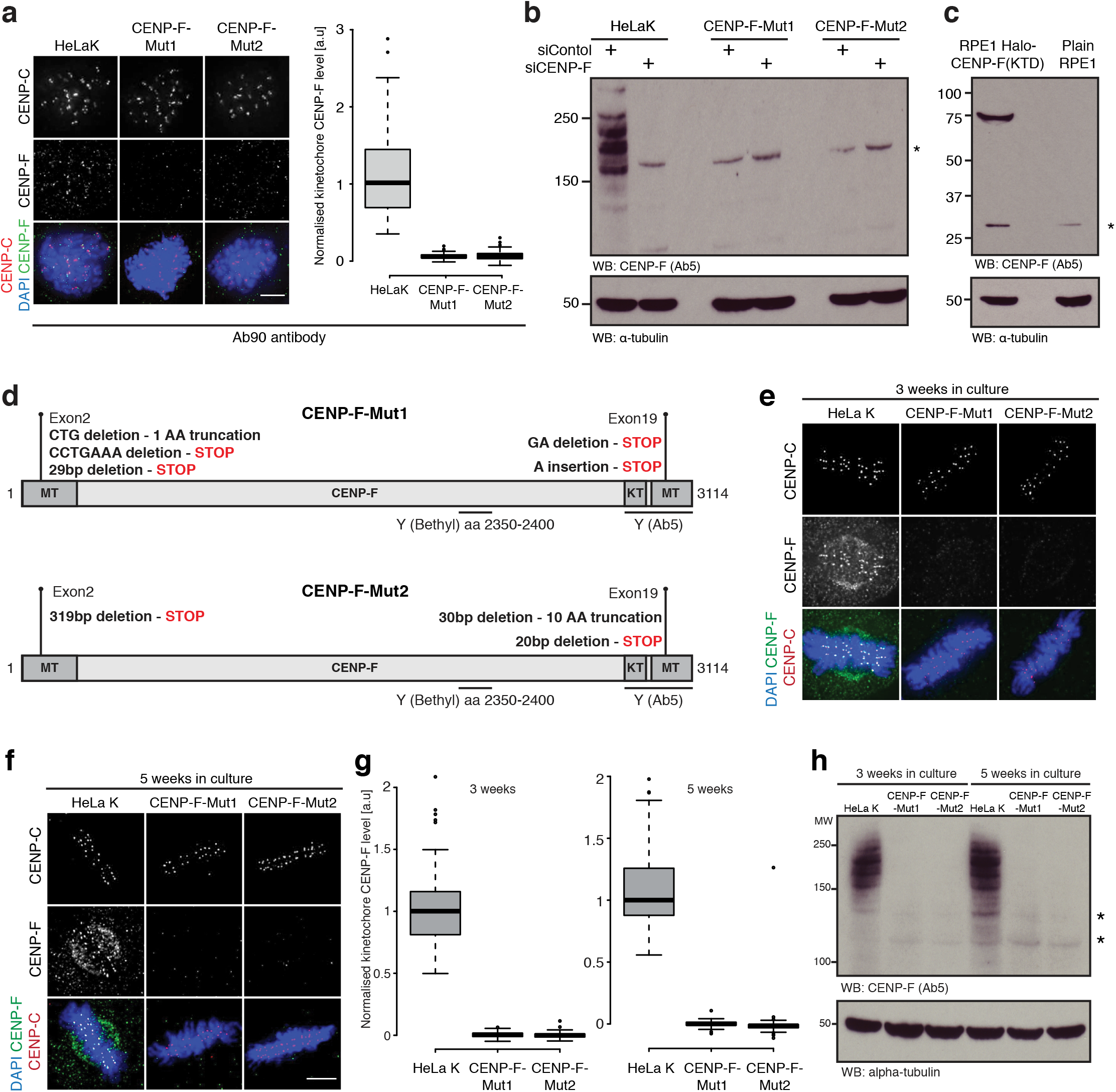
**(a)** Left Immunofluorescence microscopy images of HeLa-K, CENP-F-Mut1 and CENP-FMut2 cells stained with DAPI and antibodies against CENP-C and CENP-F(Ab90). Scale bar 5μm. Right Quantification of kinetochore CENP-F intensity relative to CENP-C in HeLa-K, CENP-F-Mut1 and CENP-F-Mut2 cells. **(b)** Immunoblot of liquid N2 protein extracts from HeLa-K, CENP-F-Mut1 and CENP-F-Mut2 cells treated with either control or CENP-F siRNA and arrested in nocodazole for 16hr. The membrane was probed with antibodies against CENP-F(Ab5) and α-tubulin. Asterisk indicates a non-specific band. **(c)** Immunoblot of liquid N2 protein extracts from Rpe1 and Rpe1-Halo-CENP-F(KTD) cells arrested in nocodazole for 16hr. The membrane was probed with antibodies against CENP-F(Ab5) and α-tubulin. Asterisk indicates a non-specific band. **(d)** Summary of CENP-F-Mut1 and CENP-F-Mut2 exon2 and exon19 sequencing. **(e)** Immunofluorescence microscopy images of HeLa-K, CENP-F-Mut1 and CENP-F-Mut2 cells cultured for three weeks and stained with DAPI and antibodies against CENP-C and CENP-F. Scale bar 5μm. **(f)** Immunofluorescence microscopy images of HeLa-K, CENP-F-Mut1 and CENP-F-Mut2 cells cultured for five weeks and stained with DAPI and antibodies against CENP-C and CENP-F. Scale bar 5μm. **(g)** Quantification of kinetochore CENP-F intensity relative to CENP-C in HeLa-K, CENP-F-Mut1 and CENP-F-Mut2 cells cultured for three (left) or five (right) weeks. **(h)** Immunoblot of liquid N2 protein extracts from HeLa-K, CENP-F-Mut1 and CENP-F-Mut2 cells grown in culture for three or five weeks. Membranes were probed with antibodies against CENP-F (Ab5) and α-tubulin.

**Supplementary figure 3.**
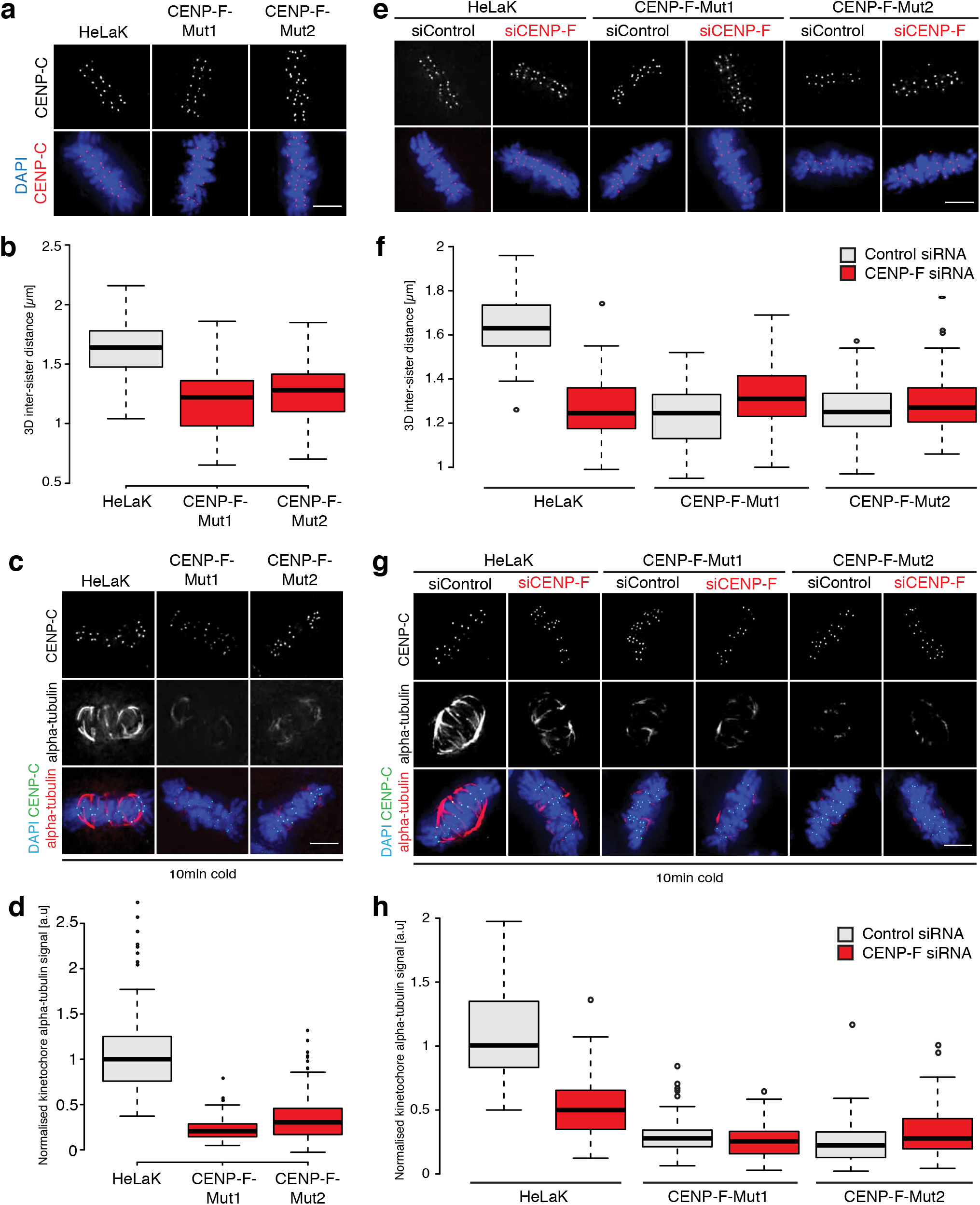
**(a)** Immunofluorescence microscopy images of HeLa-K, CENP-F-Mut1 and CENP-F-Mut2 cells stained with DAPI and an antibody against CENP-C. Scale bar 5μm. **(b)** Quantification of the CENP-C based inter-sister distance in HeLa-K, CENP-F-Mut1 and CENP-F-Mut2 cells. **(c)** Immunofluorescence microscopy images of HeLa-K, CENP-F-Mut1 and CENP-F-Mut2 cells incubated on ice for 10min before being stained with DAPI and antibodies against CENP-C and α-tubulin. Scale bar 5μm. **(d)** Quantification of kinetochore proximal α-tubulin intensity in HeLa-K, CENP-F-Mut1 and CENP-F-Mut2 cells incubated on ice for 10min. **(e)** Immunofluorescence microscopy images of HeLa-K, CENP-F-Mut1 and CENP-F-Mut2 cells treated with either control or CENP-F siRNA and stained with DAPI and an antibody against CENP-C. **(f)** Quantification of the CENP-C based inter-sister distance in HeLa-K, CENP-FMut1 and CENP-F-Mut2 cells treated with either control or CENP-F siRNA. **(g)** Immunofluorescence microscopy images of HeLa-K, CENP-F-Mut1 and CENP-F-Mut2 cells treated with control or CENP-F siRNA, incubated on ice for 10min, and stained with DAPI and antibodies against CENP-C and α-tubulin. Scale bar 5μm. **(h)** Quantification of kinetochore proximal α-tubulin intensity in HeLa-K, CENP-F-Mut1 and CENP-F-Mut2 cells treated with control or CENP-F siRNA and incubated on ice for 10min.

**Supplementary figure 4.**
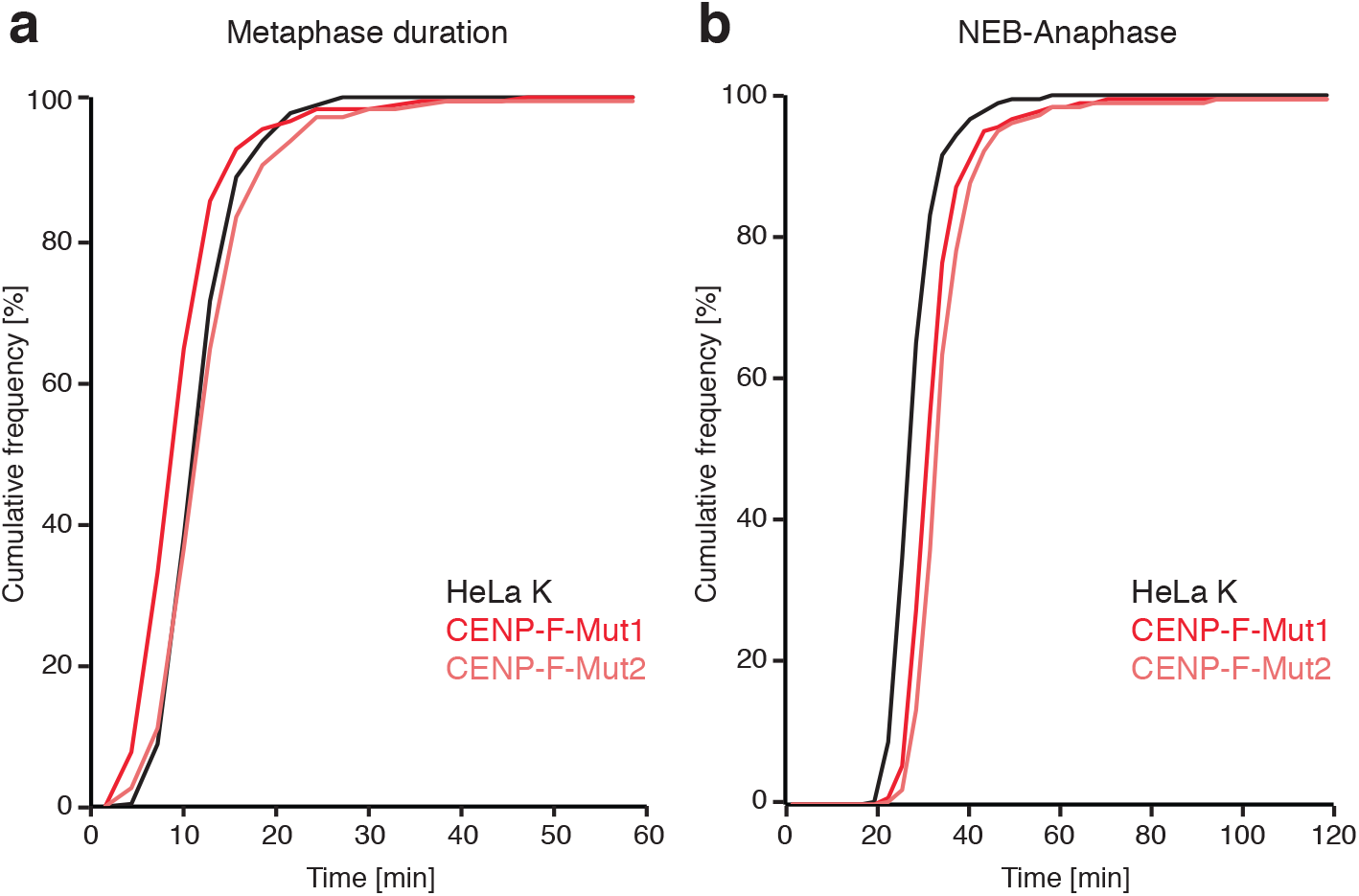
Quantification of metaphase duration **(a)** and NEB-anaphase **(b)** in HeLa-K, CENP-F-Mut1 and CENP-F-Mut2 cells. Cells were visualized using 1μM SiRDNA and imaged every 3min for 12hr.

**Supplementary figure 5.**
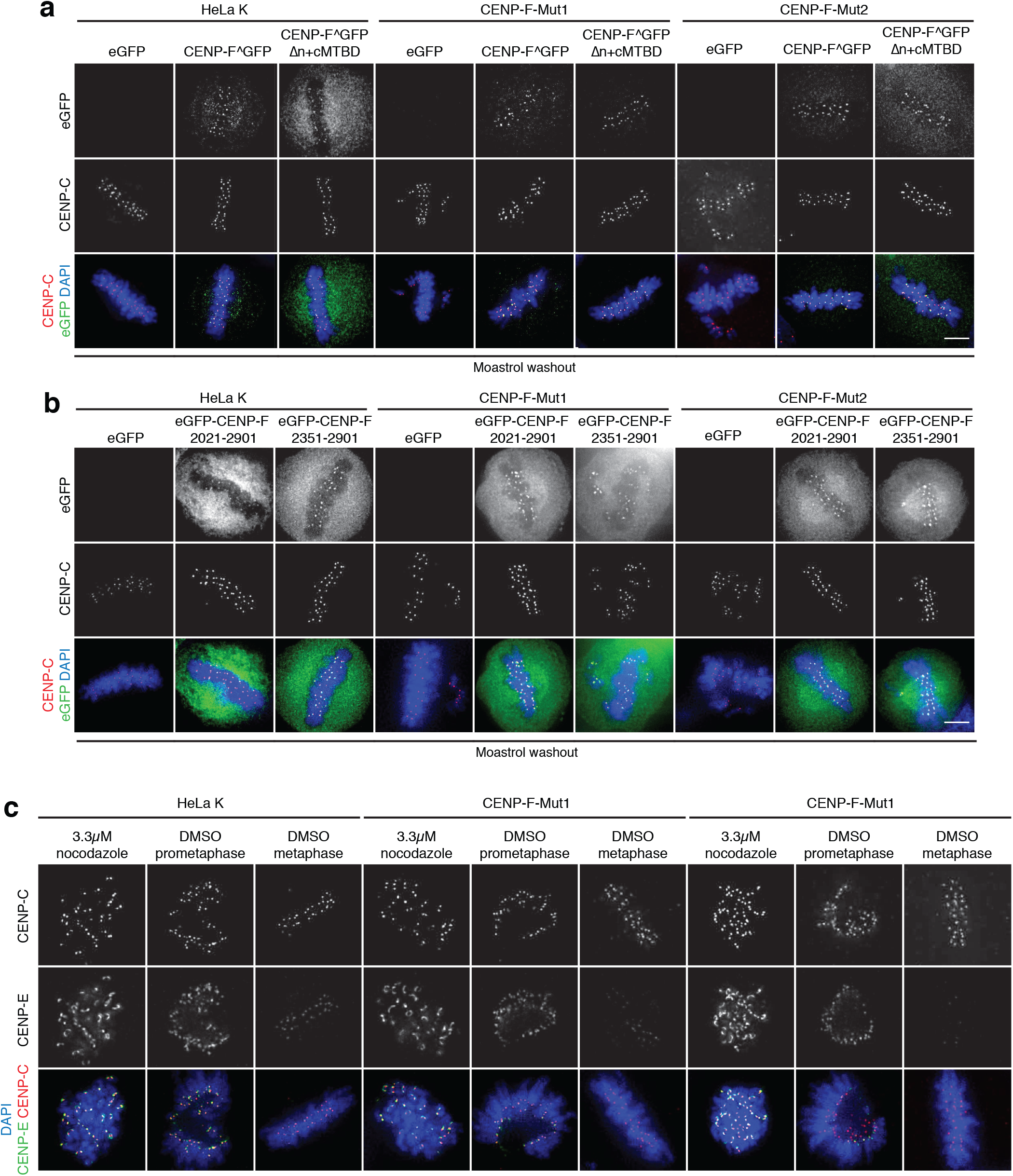
**(a)** Immunofluorescence microscopy images of HeLa-K, CENP-F-Mut1 and CENP-F-Mut2 cells transfected with eGFP, CENP-F^GFP or CENP-F^GFPΔn+cMTBD before monastrol release and staining with DAPI and an antibody against CENP-C. Scale bar 5μm. **(b)** Immunofluorescence microscopy images of HeLa-K, CENP-F-Mut1 and CENP-F-Mut2 cells transfected with eGFP, eGFP-CENP-F(2021-2901) or eGFP-CENP-F(2351-2901) before monastrol release and staining with DAPI and an antibody against CENP-C. Scale bar 5μm. **(c)** Immunofluorescence microscopy images of HeLa-K, CENP-F-Mut1 and CENP-F-Mut2 cells treated with either nocodazole or DMSO (early prometaphase and metaphase cells shown) and stained with DAPI and antibodies against CENP-C and CENP-E. Scale bar 5μm

